# A Mitochondrially Derived Plastidial Transporter Regulates Photosynthesis in the Diatom *Phaeodactylum tricornutum*

**DOI:** 10.1101/2025.02.14.638225

**Authors:** Cécile Giustini, Davide Dal Bo, Mattia Storti, Mick Van Vlierberghe, Denis Baurain, Pierre Cardol, Youjun Zhang, Alisdair R. Fernie, Duncan Fitzpatrick, Eva-Mari Aro, Guillaume Allorent, Pascal Albanese, Dimitri Tolleter, Gilles Curien, Giovanni Finazzi

## Abstract

Eukaryotic phototrophs depend on the activity of two engines (the plastid and the mitochondrion) to generate the energy required for cellular metabolism. Because of their overlapping functions, both activities must be closely coordinated. At the plastid level, optimization occurs through alternative electron transport, the diversion of excess electrons from the linear transport chain, and metabolic exchanges. A similar process takes place in the mitochondria, with documented evidence of energy and redox equivalents being exchanged between the two organelles. Organelle-organelle energy interactions at the physiological level are well established in diatoms, an ecologically significant member of phytoplankton. Yet the molecular components involved in this process remain largely unknown. Here, we identify a Mitochondrial Carrier Family (MCF) transporter, MCFc, located at the plastid envelope of *Phaeodactylum tricornutum*, which seems to be widely distributed in complex algae. We then compare the performance of a wild-type and a mutant lacking MCFc. An analysis of spectroscopic and oxygen exchange data unveiled altered energetic interactions in the mutant, suggesting that MCFc, plays a role in plastid-mitochondrion communication. In silico analysis of MCFc implies a similar substrate-specific model to that of ADP/ATP carriers, although distinct motif differences in MCFc indicate potential variations in its function, with possible substrates including arginine, aspartate/glutamate, or citrate. These findings illuminate how mitochondrial energy contributes to fueling diatom photosynthesis.

## Introduction

Photosynthesis utilises light-driven electron flow to generate bioavailable reducing power (NADPH), which is subsequently utilized by the Calvin-Benson-Bassham (CBB) cycle. Photosynthesis also requires ATP, but the ATP/NADPH ratio generated by linear electron flow is thought to be insufficient to fuel CO_2_ assimilation (reviewed in Alric 2010, and Petersen *et al*. 2012). This ratio is changeable via the modulation of linear (LEF) and alternative electron flow, comprising cyclic electron flow (CEF) around PSI (Shikanai, 2007) and the ‘water-water cycles’ (*i. e*. flavodiiron proteins and Mehler reaction). Exchange mechanisms between plastids and mitochondria have been characterised in land plants (Hoefnagel, Atkin, and Wiskich 1998). Thanks to these mechanisms, the ATP/NADPH ratio undergoes continuous adjustments according to light availability and cellular metabolic demands. In diatoms, the close proximity between plastids and mitochondria led to the hypothesis that exchanges between the two organelles occur through direct physical contacts, as shown by electron microscopy (EM) and focused ion beam milling (FIB)-EM imaging (Bailleul et al., 2015). However, recent results obtained from samples prepared in more native conditions (cryo FIB- SEM) suggest that this is unlikely the case (Uwizeye et al. 2021), reopening the question of chloroplasts-mitochondria interactions. We decided to re-evaluate the metabolic interaction mechanisms between the two organelles in diatoms, aiming at identifying specific molecular transporters. By combining molecular, bioinformatic and physiological analyses, we identified a candidate transporter (mitochondrial carrier family chloroplastic, MCF_c_) and characterised the phenotypic features of mutant lines lacking this function in the diatom *Phaeodactylum tricornutum*.

The mitochondrial carrier family (MCF) proteins are a group of nuclear-encoded and membrane-embedded proteins usually localized at the inner membranes of mitochondria. These proteins, typically small channels with a molecular mass of about 30–34 kDa, facilitate the selective transport of essential metabolites, such as di- and tricarboxylates, keto acids, amino acids, nucleotides, and coenzymes/cofactors, across the inner mitochondrial membrane, thereby linking mitochondria and the cytosol (Palmieri et al., 2011; Palmieri, 2013). This connection is vital, as many physiological processes require the cooperation of both intra- and extra mitochondrial pathways.

All MCF proteins share common structural features, consisting of three tandemly repeated homologous domains, each about 100 amino acids in length. Each domain contains two hydrophobic stretches separated by hydrophilic regions, with the signature sequence motif PX[D/E]XX[K/R]X[K/R] (PROSITE PS50920, PFAM PF00153, IPR00193), which is used to recognize MCF members (Satre et al., 2007). Many MCFs catalyse 1:1 exchange reactions (antiport), though other transport modes are also possible like uniport and H^+^-compensated anion symport (Palmieri and Pierri 2010). MCFs can be classified as either electrophoretic (electrogenic) or electroneutral based on the nature of the reactions they catalyse. For example, ADP/ATP and aspartate/glutamate carriers are electrogenic, while inorganic phosphate (Pi), glutamate, GTP/GDP, oxoglutarate, and ornithine carriers are electroneutral. Given their role in exchanging a wide variety of solutes across cellular membranes and connecting numerous metabolic pathways in different cell compartments, it is conceivable that our MCF transporter candidate belongs to this family, or at least shares some of their characteristics.

## Materials and Methods

### Phaedactylum tricornutum cultivation

The *Phaeodactylum tricornutum* (Pt1, CCAP 1055/3) cells were cultivated following the protocol outlined in Villanova et al. 2021. They were grown in a modified enriched Seawater Artificial Water (ESAW) containing a concentration of N and P ten times higher compared to the regular ESAW medium (Harrison, Waters, and Taylor 1980) modified in (Berges, Franklin, and Harrison 2001), added Cu to 3.92 10^−8^ M and removed silicate. Cells were maintained at 19°C under a light intensity of 40 μmol photon m^-2^ s^-1^ with a 12h light/12h dark photoperiod and shaking at 90 rpm. They were collected in exponential phase and concentrated to a density of 2-5 10^6^ cells per ml.

### Phylogenetic analysis

To address the origin and distribution of MCFc, we built three types of phylogenetic trees, all derived from previously published large-scale datasets (Van Vlierberghe et al. 2021a-b) with progressively less outgroup sequences included. First, Supplementary Figure 1A shows the original tree computed from one of the MAFFT-aligned (Katoh & Standley 2013) broadly sampled orthogroups (OG0001460) obtained in Van Vlierberghe et al. (2021a). Second, Supplementary Figure 1B shows its largest photosynthetic clan (= subtree) (Van Vlierberghe et al. 2021a), further enriched (using Forty-Two [Irisarri et al. 2017]) with sequences from 172 high-quality transcriptomes of plastid-bearing lineages resulting from the decontamination, pooling and dereplication of the 678 MMETSP samples (Keeling et al. 2014) performed in Van Vlierberghe et al. 2021b. The third one, Supplementary Figure 1C (and Supplementary Figure 1B) is focused on MCFc itself. Amino-acid alignments were filtered with ali2phylip.pl from the Bio::MUST::Core software package available on CPAN [https://metacpan.org/dist/Bio-MUST-Core] and used to infer phylogenetic trees using IQ-TREE 2 (Minh et al. 2020) with the ultrafast bootstrap option under the C20 (Le et al. 2008) (or LG4X [Le et al. 2012]) model. Raw trees were then automatically rooted, annotated, colorized and collapsed based on NCBI Taxonomy using format-tree.pl (also from Bio::MUST::Core) and uploaded to ITOL for further annotation and graphical rendering (Letunic et al. 2024). Alignments and trees are available at Figshare under DOI:10.6084/m9.figshare.28207955.

### Localization

The construction of the MCFc::GFP fusion was performed by Liu et al., 2025. To visualize mitochondria, the mitochondrial probe TMRM (tetramethylrhodamine, methyl ester, perchlorate, Invitrogen MitoProbe) was added at a concentration of 100 nM. Imaging was performed using a Zeiss LSM 900 confocal microscope equipped with ZEN software (version 3.0, Blue edition). A 63x/1.4 M27 objective lens was used in confocal mode with an Airyscan 2 detector to ensure fast acquisition without compromising sensitivity, aided by a collimation optic. Excitation for chlorophyll fluorescence was provided by a 405 nm blue laser, and emission was collected between 650 and 700 nm. GFP visualization involved 488 nm excitation with fluorescence collected between 500 and 536 nm, while for TMRM, excitation was at 561 nm with fluorescence collected between 570 and 613 nm.

### Construction of the MCFc mutants

*Phaeodactylum tricornutum* mutant MCFc strains have been generated through CRISPR/Cas9 according to the protocol described in Giustini *et al*. 2024.

The following target sequence was designed using the PhytoCRISP-Ex web tool (Rastogi et al. 2016):

MCF2: 5’-GTCAGCCGAGAAGCCGACGA-3’;

The target sequences were inserted into the pKSdiaCas9_sgRNA plasmid (Addgene #74923). The resulting plasmids and the pAF6 plasmid, conferring a resistance to zeocin (100 μg/mL) were coated on tungsten beads and used for biolistic transformation of the cells with the PDS- 100/He system (Bio-Rad, Hercules, CA, USA; 1672257). Genomic DNA from zeocin resistant transformants was extracted, amplified by PCR using the following primers 5’- ACGCGCATGTAGTTACAGTTAGCGTATTT-3’ (forward) and 5’-CCTGGGTAGCGGGAAGCGTTT-3’ (reverse) and sequenced for mutant identification.

### Protein extraction and immunoblot analysis

Proteins were extracted and analyzed as described in Seydoux et al. 2022. In short, cells (approximately 8.0×10^6^) were pelleted and resuspended in 50 mM HEPES buffer (pH 7.5) supplemented with an EDTA-free protease inhibitor cocktail (cOmplete, Roche). Cell lysis was achieved using a Precellys Evolution Homogenizer (Bertin, France) with the following settings: 2 cycles of 30 seconds at 10,000 rpm, with a 30-second pause between cycles at 4°C. Proteins were then precipitated using 80% acetone by centrifugation at 4°C and resuspended in lysis buffer (100 mM Tris-HCl, pH 6.8, 4% SDS, 20 mM EDTA) supplemented with the protease inhibitor.

For immunoblot analysis, 10-30 μg of protein were separated by SDS-PAGE in Tris-Glycine buffer (25 mM Tris, 190 mM glycine, 0.05% SDS). Proteins were transferred onto a nitrocellulose membrane using the same buffer supplemented with 20% ethanol for 80 minutes at 100 V. Immunodetection was performed using a guinea pig anti-MCFc serum raised against the full-length recombinant protein (dilution 1:3,000; Charles River Laboratories, Strasbourg, France) and an anti-ATPB antibody (dilution 1:5,000; AS05085; Agrisera). Detection was carried out with an HRP-conjugated secondary antibody, and the signal was developed using the Clarity Western ECL Substrates kit. Images of the blots were captured using a CCD imager (ChemiDoc MP Imaging System, Bio-Rad).

### Oxygen exchange measurements

Oxygen consumption (respiration) and production (photosystem II) were monitored using oxygen electrode systems that were temperature-controlled at 19°C (Hansatech Instruments, UK).

Measurements with stable isotopes of carbon (^13^C) and oxygen (^18^O) in MIMS were performed as described in Fitzpatrick et al. 2022. Cells were taken at around 2.10^6^ cells. mL^-1^ and concentrated to approximately 10μg chlorophyll. ml^-1^ in darkness. The sample was purged with N_2_ to minimize background ^16^O_2_ before a bubble of ^18^O_2_ (99% Cambridge Isotope Laboratories Inc, UK) was loaded into the stirring liquid, bringing the concentration of the heavier isotope up to approximately 500 μM. ^13^C bicarbonate was added to 10mM concentration. The conversion to ^13^CO_2_ was enhanced by adding a small volume of Carbonic Anhydrase (Sigma, USA) at 1 mg. mL^-1^. Data were recorded in darkness and later at 25 and 250 μmol photons m^-2^. s^-1^. Measurements were performed for 5 minutes at each light condition. All data were analyzed, and fluxes were calculated with equations described in Beckmann et al. (2009), which include offsets for the changing relative concentrations of ^16^O_2_ and ^18^O_2_.

### Fluorescence and spectroscopic characterization of photosynthetic light reactions

Chlorophyll fluorescence was imaged in 100 μL of cell suspension placed on a 96-well plate. Cells were dark-adapted for 10 min before measurement. Fluorescence was quantified using a Speedzen III fluorescence imaging setup (JBeam Bio, France). Maximum fluorescence in the dark (F_m_) or during actinic light exposure (F_m’_, 900 μmol photons m^-2^ s^-1^) was measured using saturating red pulses (250 ms, 3’000 μmol photons m^-2^ s^-1^) followed by blue light (470 nm) detection pulses (10 μs). The chlorophyll fluorescence was then measured before (F_0_) and during the saturating pulse (F_m_), and the Electron Transfer Rate (ETR) was calculated as (F_m_’- F)/F_m_’ x PAR x 0.5, where PAR is the intensity of actinic light used to stimulate the photosynthesis and the term 0.5 represents the 50% of probability that one electron goes to the PSII (instead of the PSI).

In vivo spectroscopic analysis was performed with a JTS-10 spectrophotometer (Biologic, France), equipped with a white LED source filtered through appropriate interference filters, and BG 39 filters to protect the photodiodes from actinic light. This light was used as a measuring beam, while actinic light induced photosynthetic change, including changes in the pmf revealed by ECS changes measured at 520-545 nm. This difference disentangles ECS signatures from spurious changes associated with either cytochrome b6f complex activity (Joliot & Joliot, 1994) or with the 535 nm shift that is linked to reflects a modification of the xanthophyll zeaxanthin during the onset (or the relaxation) of NPQ (Johnson & Ruban, 2014) The amplitude of the ECS signal was normalized to a signal corresponding to 1 charge separation, i.e. the amplitude induced 150 μs after exposure to saturating single turnover laser flashes. Because PSII was inactivated by the addition of saturating amounts of the inhibitors DCMU (20 μM) and hydroxylamine (1 mM), only PSI is active in these conditions, and therefore, the ECS amplitude corresponds to one charge separation per photosynthetic electron transfer chain (Bailleul et al. 2010).

### Metabolite analysis

Metabolites were extracted and analyzed as described in Villanova et al. 2017. Ten million cells were harvested on a Durapore-HV membrane filter disc 2.5 cm in diameter and with a pore size of 0.45 μm (Millipore, USA) by vacuum filtration. The filter with the cells was then transferred into a 1.5 ml tube and frozen in liquid nitrogen. Frozen samples were stored at −80°C until metabolite extraction. Metabolites were extracted by immersing the filter in 1 ml of 90% (v/v) methanol containing 0.1 μg ml^−1^ U ^13^C sorbitol as an internal standard. The tubes were sonicated in a water bath-type sonicator for 1 min in ice cold water and then incubated at 4°C for 1 h with shaking. The remaining solution was centrifuged at 22 000*g* for 5 min at 4°C. A 50 μl aliquot of the supernatant was used to determine chlorophyll concentration, while a 900 μl aliquot was reduced to dryness using a SpeedVac vacuum concentrator (Thermo Fisher Scientific, USA). Dried samples were stored at −80°C after filling the tubes with argon gas. The metabolite profile was determined exactly as described in Obata et al. 2013.

### Statistical analysis

For all statistical analyses, excluding phylogenetic analysis, unpaired t-tests with Welch’s correction were performed using GraphPad Prism (version 10.4.1 for Mac OS X, GraphPad Software, Boston, Massachusetts, USA, www.graphpad.com). Statistical significance was denoted as follows: *p* < 0.05 (\**), p < 0*.*01 (\*\**), and *p* < 0.001 (***)

## Results

### Identification of a putative transporter involved in chloroplast-mitochondria energetic interactions

While functional analysis of diatoms supports the existence of energy exchanges between plastids and mitochondria, the specific molecular candidates driving this process remain unidentified. To address this question, we searched among proteins with putative complex plastid target sequences based on ASAFind (Gruber et al., 2015), a tool designed to identify nuclear-encoded plastid-localized proteins in the red algae lineage. This tool relies on the output from the SignalP web tool (Petersen et al., 2011). It recognizes conserved “ASAFAP” motifs (for transit through the ER membrane surrounding the plastid in diatoms) as well as transit peptides. Among candidates, we focused on homologs of transporters involved in ATP or reducing equivalent transport and selected a putative MCF transporter, MCFc (Phatr3_J46742, also PTI_11G01630 in Pico-PLAZA 3 and PTI_11G01280 in Pico-PLAZA 2). According to the Ensembl Protists Gene Browser (protists.ensembl.org), Phatr3_J46742 is located on Chromosome 11 and encodes the protein B5Y3N0 (UniProt). The predicted structure of MCFc has been modeled with high confidence in the AlphaFold database (Varadi et al. 2024) (Figure 1A, Supplementary Figure 1). We performed a 3D structural homology search using the PDB repository and AlphaFold DB (van Kempen et al. 2024), revealing clear similarities to four well-characterized mitochondrial transporters in mammals and fungi. This approach, along with its classification within the MCF family, suggests that MCFc transports amino acids, carboxylic acids, fatty acids, cofactors, inorganic ions, or nucleotides (Ruprecht and Kunji 2020). Based on its structural similarity to known plastid-targeted MCF transporters such as the land plant ADNT1 for ADP/ATP exchange and an arginine transporter (BAC1), both from Arabidopsis thaliana (Supplementary Table 1, Supplementary Figures 2 and 3), MCFc appears to be a plastidial membrane protein that potentially plays a key role in chloroplast- mitochondria crosstalk.

**Figure 1:**
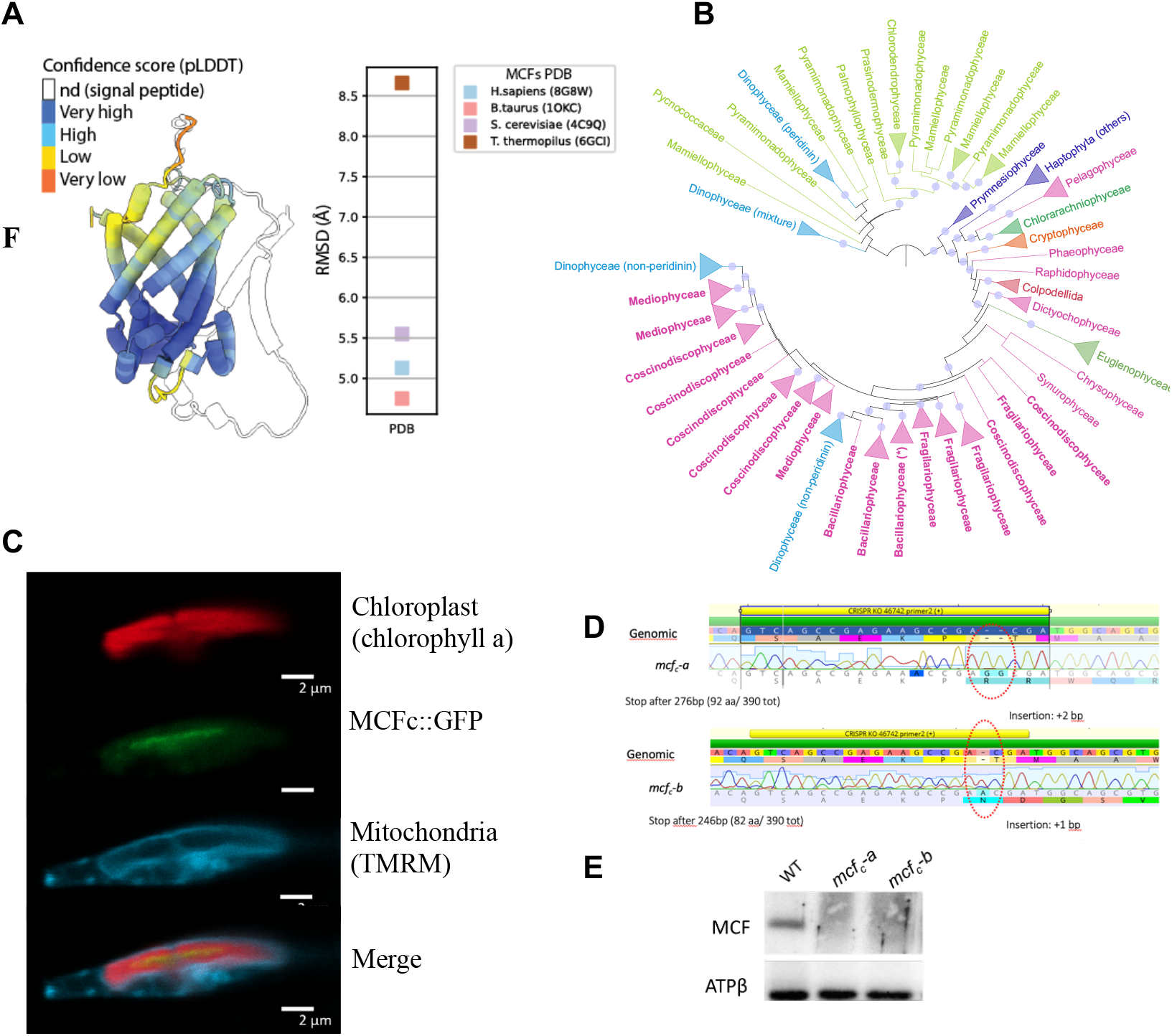
MCFc is a plastidial member of the Mitochondrial Carriers Family. **(A)** Predicted structure of MCFc and the Root mean square deviation of atomic positions in Angstroms (RMSD) for each amino acid backbone αCarbon is indicated from canonic MCF **(B)** Phylogenetic tree of MCFc inferred under the C20 model. Monophyletic groups are collapsed at the family level and colored by taxonomic affiliation (light green: green algae, darker greens: chlorarachniophytes and euglenids, orange: cryptophytes, violet: haptophytes, pink: ochrophytes, blue: dinophytes, red: colpodellids). Diatoms are shown in boldface and the group including *Phaeodactylum* MCFc (Phatr3_J46742) is denoted by an asterisk (*). Nodes with ultrafast bootstrap support (UFBS) ≥ 85% are indicated by a semi-transparent circle. **(C)** Subcellular localization of MCFc. Fusion MCFc_GFP shows a colocalization with the chlorophyll fluorescence and not with mitochondria. **(D)** Molecular characterization of MCFc KO mutants. Sequencing of the Mcf c gene in mcf-a and mcf-b indicates that both clones bear a stop codon after 90 codon triplets due to the insertion of 2 and 1 base pair, respectively. **(E)** Western blot analysis confirms the successful knocking out of the Pt-MCF gene by revealing the absence of a specific protein at around 40 KDa in the two mutant lines, ATPβ is the loading control.

Homologs of MCFc were analyzed in a dataset generated from orthologous groups inferred from high-quality proteomes (Van Vlierberghe et al. 2021). The corresponding orthogroup (OG0001460) included three sequences from *Phaeodactylum tricornutum*. A preliminary phylogenetic tree (C20 model, Supplementary Fig. 1A) showed that one gene copy (PTI_10G03480 = Phatr3_J46612, putative mitochondrial copy) belongs to a large group containing both photosynthetic (PS) and non-PS lineages (UFBP support 76%). Another copy (PTI_01G05580 = Phatr3_J42874) is specific to a group limited to PS heterokonts (= Ochrophytes) (UFBS 89%). In contrast, the third copy (PTI_11G01280 = Phatr3_J46742 = MCFc) is in a large group that includes most PS lineages, among which non-streptophyte green algae, to the notable exception of red algae (UFBS 70%). Finally, many diatoms (but not *Phaeodactylum*) and a few other lineages display a fourth copy that was provisionally considered as mitochondrial too (UFBS 55%).

The two PS subtrees (OG0001460-8) were enriched with high-quality sequences from PS lineages (Van Vlierberghe et al. 2021b) to clarify the phylogenetic environment of the diatom MCFc gene, yielding a relatively well-supported tree (C20 model, Supplementary Figure 1B). The absence of red algae suggests they have lost this gene copy entirely. Ochrophytes are split into two distinct groups, corresponding to the two remaining gene copies of *Phaeodactylum*, with diatoms each time forming a maximally supported monophyletic group (UFBS 96-100%), thus confirming the usefulness of this organism as a model diatom in the present work. Similarly, haptophytes are split into two (unequal) groups, but both coincide with the gene copy in most PS lineages.

In the large PS subtree, cryptophytes non-robustly (UFBS 25-55%) appear near the base of a densely sampled group of red complex algae (i.e., ochrophytes, colpodellids), thus likely exhibiting a “red copy,” even though the highly supported (UFBS >90%) nested positions of complex green euglenids and chlorarachniophytes may seem odd. Surprisingly, most haptophytes group with green algae rather than with this red copy. Moreover, the positions of dinophytes show strong evidence (UFBS >90%) of lateral gene transfer (LGT) from either ochrophytes (non-peridinin plastids) or more strangely from green plants (both peridinin and non-peridinin plastids), suggesting a late replacement of the red copy in dinophytes. This analysis used the ochrophyte-specific copy as the outgroup to root the MCFc tree. A third tree focusing on the MCFc subtree alone was built to further analyze those relationships after pruning the ochrophyte-specific gene copy. In this well supported tree (Supplementary Figure 1C and Fig. 1B), green algae were used as the outgroup, which is coherent with the relatively long branch (UFBS 99%) separating their “green copy” from the red copy of all complex algae. Nonetheless, this new tree identifies MCFc as an additional example of a gene with a convoluted history in PS lineages, especially those of complex algae (Petersen et al. 2014).

Subcellular localization studies using a MCFc::GFP fusion showed co-localization with the chlorophyll fluorescence signal, especially at the periphery of the chloroplast (probably at the plastid envelope) and not with mitochondria (Figure 1C, see also Figure 3 from Liu et al. 2025). The subcellular localization is, therefore, consistent with the hypothesis that this MCF member belongs to a peculiar subfamily only present in PS organisms, except the Streptophyta and Rodophyta.

### Generation of MCFc Knock-Out Mutant Strains

To investigate the potential role of MCFc in chloroplast-mitochondria interactions, we generated knockout (KO) strains using CRISPR-Cas9 technology (Giustini et al., 2024). After transformation, zeocin-resistant *P. tricornutum* (Pt1) colonies were sequenced, revealing two independent clones with mutations in the Phatr3_J46742 gene (MCFc). These mutations introduced a stop codon after 90 or 82 amino acids due to the insertion of 2 and 1 base pair(s), respectively (Figure 1D).

We confirmed the absence of the MCFc protein by immunodetection analysis using a custom-made antibody that recognized a 40 kDa band in the wild type (WT), corresponding to the predicted molecular weight of the MCFc protein. This band was absent in the mutant lines, confirming the successful generation of knockout lines for the Phatr3_J46742 gene (Figure 1E).

To explore the potential role of MCFc in diatom energetics, we examined the growth and photosynthetic capacity of the two knockout (KO) mutant strains. We combined the results from both clones because their phenotypes were indistinguishable. Although no growth defects were observed (Figure 2A), we found that the KO lines had lower photosynthetic performance (Electron Transport Rate - ETR) compared to wild-type (WT) cells (Figure 2B). To complete the physiological characterization of the mutant, we assessed its ability to maintain a proton motive force (ΔΨd) in the dark, a functional trait linked to chloroplast-mitochondria interactions (Bailleul et al., 2015). This parameter was quantified using the Electrochromic Shift (ECS) signal, which responds to changes in membrane ΔΨ with linear (at 520 nm) and quadratic (at 565 nm) dependencies (Bailleul et al., 2015; Joliot & Joliot, 1989). The ΔΨd signal was approximately 25% lower in the mutant strains compared to the WT (Figure 2C), indicating that MCFc contributes to energy interactions between these two organelles. However, this role appears to be partial and is likely complemented or compensated by other metabolic exchanges, which may help to mitigate the impact of MCFc disruption.

**Figure 2:**
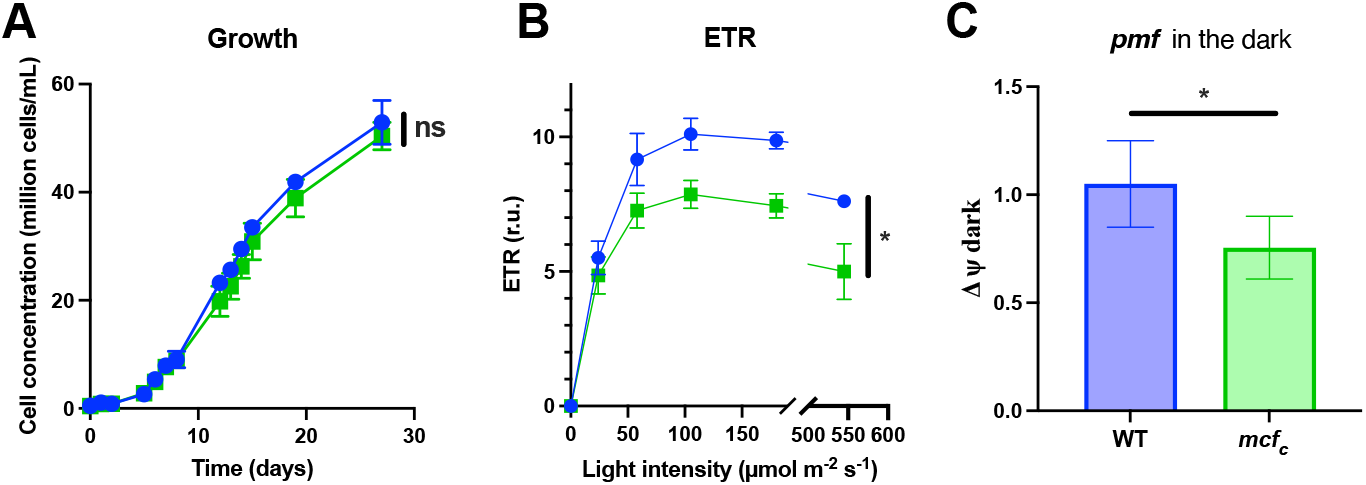
MCFc impacts photosynthesis with no consequence on growth. In blue the WT, in green *mcf*_*c*_ **(A)** Growth of *Phaeodactylum tricornicum* in autotrophic condition with moderate light regime (12h/12h light cycle, 50 μmol photon m^-2^ s^-1^) has been done for the WT and mutant *mcf*_*c*_. Data ± SD from 6 biological replicates. **(B)** Photosynthetic efficiency (plotted as ETR) has been measured by chlorophyll fluorescence at different light intensities for the WT and *mcf*_*c*_. Data ± SD from 6 biological replicates. **(C)** The Proton Motive Force (*pmf*) and specially his ΔΨ component have been calculated using the Electro Chromic Shift (ECS) signal measurements. Data ± SD from 6 biological replicates. See methods for more details.

### MCF_c_ modulates the relationship between photosynthesis and respiration

We monitored the relationship between oxygen production (from water-splitting PSII) and oxygen consumption (via respiration) (Figure 3A). In diatoms, these processes are linearly correlated, suggesting the existence of a link between the export of excess reducing power from the plastid to the mitochondria, its use for ATP production, and the subsequent import of ATP into the chloroplast to drive CO_2_ assimilation (Bailleul et al., 2015). Since MCFc may influence photosynthetic electron transfer rates (ETR) and the membrane potential in the dark (ΔΨd) as part of the proton motive force, we examined the interplay between photosynthesis and respiration under varying concentrations of SHAM and Antimycin A, two inhibitors of mitochondrial respiration that gradually reduce mitochondrial activity. We observed a linear relationship between the two mechanisms (respiration and photosynthesis) in WT cells (Figure 3A), indicating that increased activity in one process enhances the other. This result aligns with previous findings (Bailleul et al., 2015). The linearity observed here was maintained in the absence of MCFc, although the slope of oxygen evolution versus consumption was less steep (Figure 3A), suggesting that organelle crosstalk was at least somewhat disrupted.

**Figure 3:**
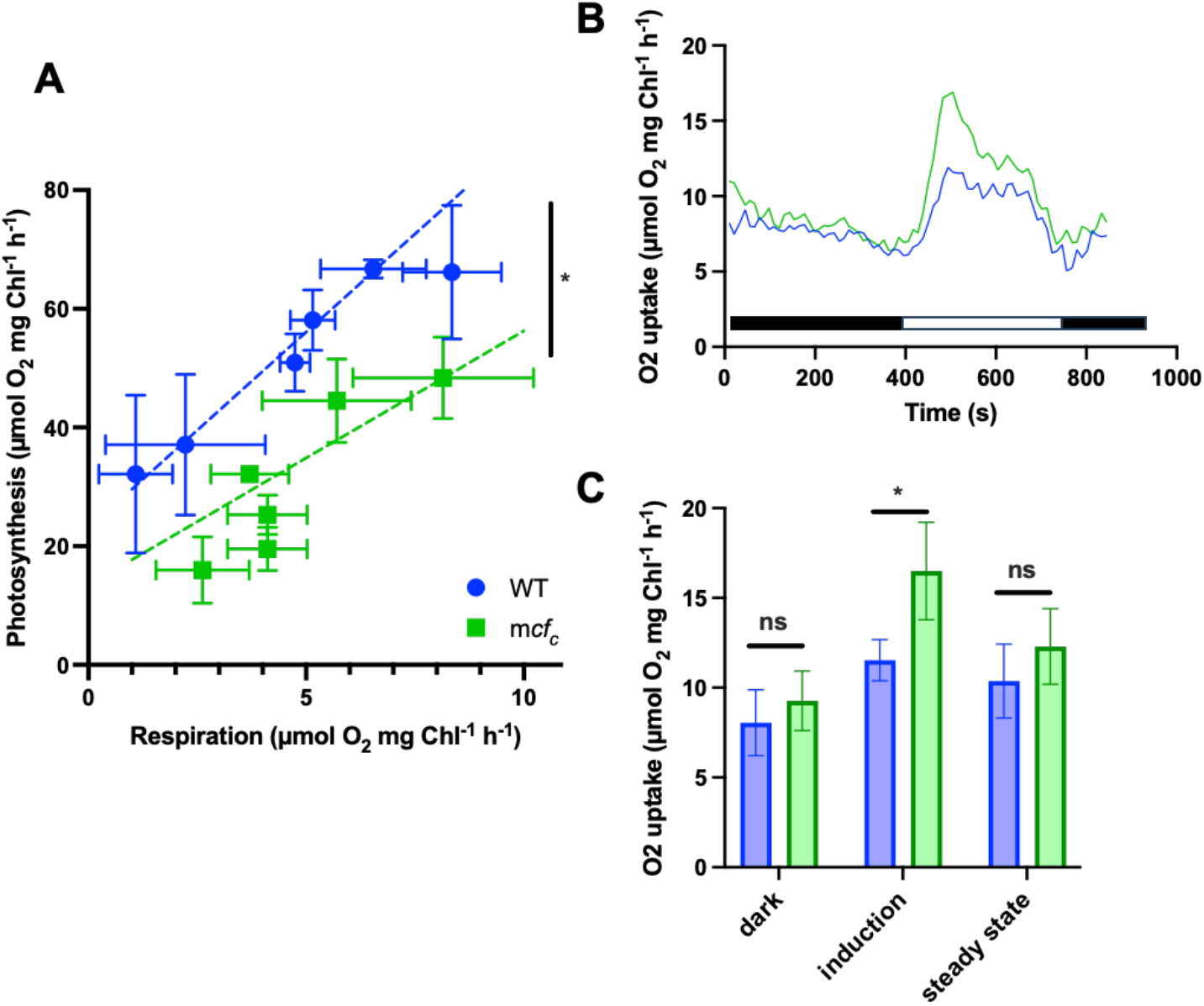
Analysis of the photosynthesis/respiration relationship. In blue the WT, in green *mcf*_*c*_ **(A)** In *P. tricornutum* cells collected in exponential growth phase, photosynthetic activity was measured as O_2_ evolution (in the light) while respiration is the rate as O_2_ uptake in the dark. Data±SD from 6 biological replicates. **(B) and (C)** O_2_ uptake rates measured by Membrane Inlet Mass Spectrometer (MIMS) in presence of ^18^O-labeled O_2_. Here is a representative trace. Cells were dark adapted before the experiment. Black bars indicate the dark period, white bar the light period **(C)** Average +/- SD of 4 independent replicates extracted from B)

We corroborated these findings using the more specific Membrane Inlet Mass Spectrometry (MIMS) technique (reviewed in Burlacot et al., 2020). MIMS is particularly well-suited for detecting dynamic, short-term effects, such as alternative flows through less efficient transporters driven by elevated substrate concentrations, often obscured under steady-state conditions by compensatory mechanisms. By employing ^18^O and ^13^C isotopes, we simultaneously recorded photosystem (PS) II O_2_ evolution and cellular respiration rates under varying light conditions. Photosynthetic O_2_ evolution and ^13^CO_2_ fixation rates followed a similar trend in the mutant and WT strains, showing an approximately linear relationship with light intensity. However, the ^18^O_2_ uptake rate increased sharply under high-light conditions in the mutant (Figure 3B, C).

Additionally, the oxygen uptake of the *mcf*_*c*_ strain exhibited a brief spike (lasting about 10 seconds) after transitioning to high light before stabilizing at a lower steady-state uptake, suggesting a transient accumulation of undefined respiratory substrates due to the absence of the MCFc transporter. Overall, the O_2_ evolution-to-uptake ratio was significantly higher in the *mcf*_*c*_ mutant compared to the WT under high light conditions (Figure 3B, C), explaining the differential slope observed in Figure 3A. Based on the notion that MCFc is involved in energy exchanges between the chloroplast and mitochondria, this transient increase in O_2_ consumption might indicate increased respiration activity due to an over-reduction of PSII.

### Metabolite analysis-based analysis of chloroplast-mitochondria exchanges

The O_2_ burst described above might illustrate a limitation on the export of metabolites between the two organelles. We performed a comparative metabolomics analysis by MCFc to investigate the nature of this metabolite (these metabolites). The absence of MCFc resulted in a significant decrease in several amino acids, including aspartic acid (Asp), glutamic acid (Glu), asparagine (Asn), leucine (Leu), and 2-ketoglutarate (2-KG, the precursor of Glu). In contrast, arginine (Arg) levels were elevated in the mutant compared to the WT (Fig. 4).

**Figure 4:**
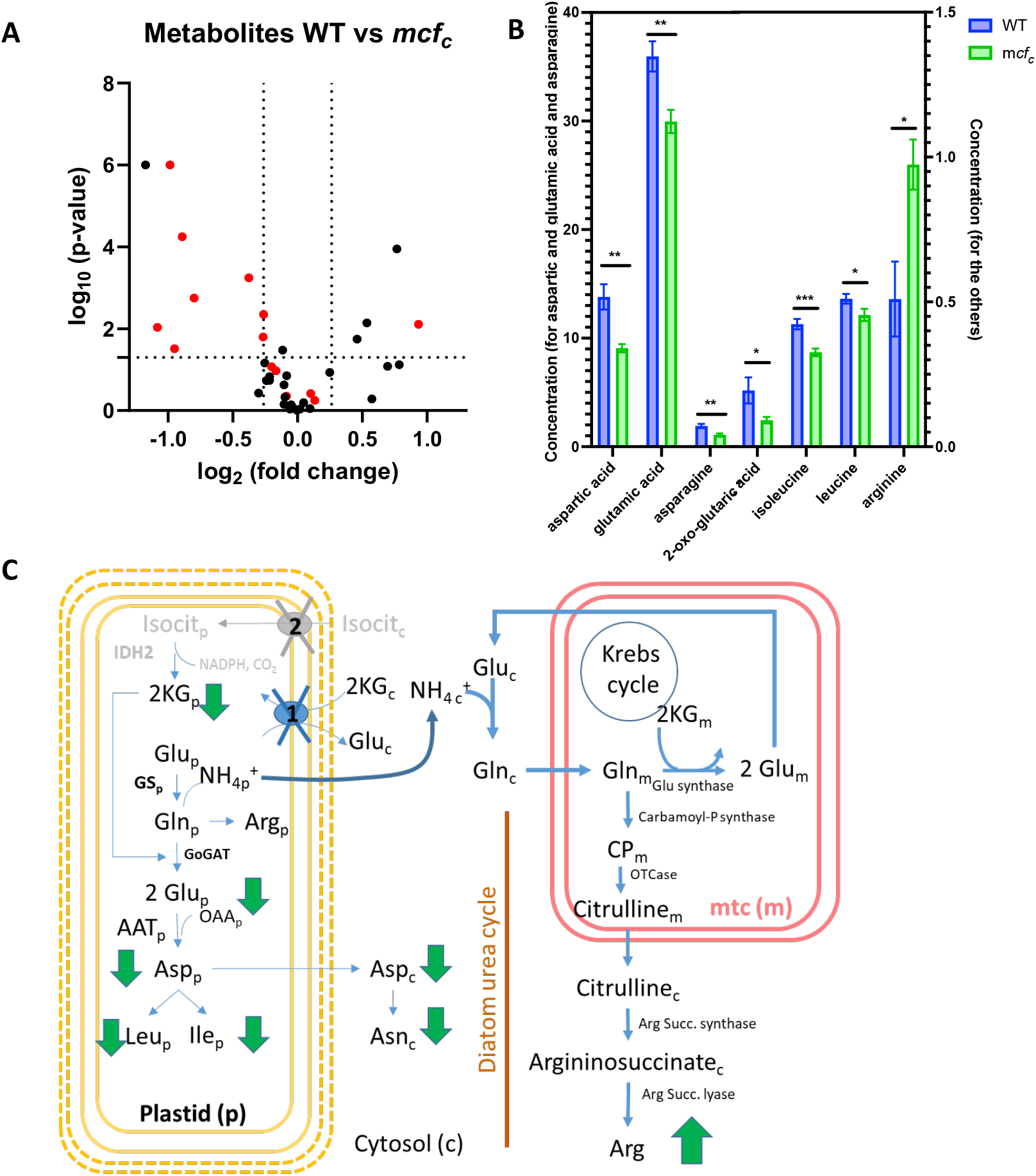
Metabolic flexibility. **(A)** Volcano plot showing the differential concentration of metabolites in *mcfC* relative to the *WT*. (the amino acids group is in red) **(B)** amino acids concentration in WT and *mcf*_*c*_. Data ± SD from minimum 6 biological replicates. **(C)** Schematic representation of metabolite changes in *P. tricornutum* cells. Flux changes upon mutagenesis are highlighted.

We interpret these changes as follows (Fig. 4C): The reduced levels of Glu and 2-KG likely originate from the chloroplast, which is about 10 times larger than the mitochondrion (Uwizeye et al., 2021). Usually, nitrogen fixation occurs in the plastid through the combined action of glutamine synthetase and GOGAT, but *de novo* 2-KG synthesis happens exclusively in the mitochondrion. Therefore, our first hypothesis would be that 2-KG import into the plastid of the MCF mutant is hindered, which could account for the decreased Glu synthesis. Since Glu is required for Asp production, a reduction in Glu would lower Asp concentrations in the plastid [AspP in Fig. 4C]-assuming amino acid demand is maintained and subsequently reduces amino acids derived from Asp, such as Leu, isoleucine (Ile), and Asn (following AspP export).

As a result, nitrogen fixation in the MCF mutant may rely more heavily on cytosolic glutamine synthase, given the reduced utilization of NH_4_^+^ in the plastid due to decreased 2-KG availability and the diffusion of NH_4_^+^ from the plastid. The interplay between nitrogen and carbon metabolism indicates that carbon is essential for nitrogen assimilation and vice versa (Smith et al. 2019; Obata et al. 2013; Huppe et Turpin 1994). Consequently, excess nitrogen in the cytosol could be redirected toward Arg synthesis via cytosolic glutamine synthase and potentially toward a hypothesized urea cycle in diatoms (Allen, 2011), which also involves arginine (as shown in the lower right of the scheme). These adjustments align with the need for nitrogen to sustain photosynthetic processes while maintaining carbon assimilation. Overall, these changes explain the significant increase in cytosolic Arg, the most prominent difference observed among the metabolites analyzed (Fig. 4C, hypothesis 1).

Alternatively, deficient import of isocitrate into the plastid (Fig. 4C, grey pathways) could also lead to a similar metabolic phenotype (hypothesis 2), as isocitrate could provide 2-KG via plastidial isocitrate dehydrogenase (ICDH) activity (Huang et al., 2021) (see Discussion).

### In-silico analysis of MCFc putative structure and substrate specificity

The transport mechanism of mitochondrial ADP/ATP carriers is well-described and likely generalizable to other mitochondrial carriers (Ruprecht and Kunji, 2020; Palmieri and Pierri, 2010). These carriers typically have a single substrate-binding site and two substrate recognition gates on each side of the membrane (Supplementary Figure 4). The coordinated movement of six transmembrane (TM) helices facilitates the alternating opening and closing of the matrix or intermembrane space (confluent with the cytosol), preventing proton leakage from the lumen. The coupling between TM helices 1-2, 3-4, and 5-6 is driven by interactions between similar sequence motifs that complement the joint TMs within the doublets, conferring substrate specificity (Ruprecht and Kunji, 2020).

An analysis of the corresponding motifs in MCFc homologs, based on structural alignment and motif scoring (Supplementary Figure 4), reveals that the odd-numbered TM helices (1, 3, and 5) share very similar motifs. At the same time, this pattern does not hold for the even-numbered TMs, suggesting a potentially different structural mechanism. By evaluating MCFc similarity to known MCF proteins, possible substrates could include aspartate/glutamate, ATP- Mg/Pi, or citrate (Supplementary Table 1).

## Discussion

In this work we searched for potential factors involved in optimizing photosynthesis in diatoms. These organisms, which are representatives of the complex red lineage phototrophs, share the most fundamental characteristics of oxygenic photosynthesis in primary endosymbionts. However, a complete understanding of the molecular components responsible for optimizing the photosynthesis process in diatoms remains incomplete.

### Optimising ATP/NADPH stoichiometry and strategies

Photosynthesis relies on maintaining an optimal ATP/NADPH ratio, which is established during the light reactions. This balance is typically regulated by adjusting the contributions of linear and alternative electron flow processes. One mechanism involves the export of reducing equivalents from the plastid to mitochondrial oxidases (Kinoshita et al., 2011). This process enables mitochondria to support photosynthetic activity under specific conditions and has been observed in green plants, with particularly high activity in some microalgae.

For instance, knockout strains with double mutations in respiratory complexes (Cardol et al., 2009; Lemaire et al., 1988) exhibit severe growth defects underscoring the critical role of mitochondrial function in photosynthetic organisms in *Chlamydomonas*. Remarkably, the *Chlamydomonas* mutant strain Fud50su, which lacks the plastidial ATP synthase, can restore phototrophic growth in a revertant strain. This recovery is achieved through enhanced ATP import from mitochondria via the ATP/ADP transporter (Merchant et al., 2007; Lemaire et al., 1988), highlighting the interplay between plastids and mitochondria energy metabolism.

In diatoms, direct energetic exchange plays a crucial role (Bailleul et al., 2015; Murik et al., 2019). Computational simulations and experimental analyses support that organelle crosstalk can regulate light-dependent metabolic changes in these organisms (Prihoda et al. 2012). In addition to direct metabolic exchanges, “indirect” interactions are also well documented: CO_2_ produced from oxidative phosphorylation can be directly fixed in the plastid (Busch et al., 2013; Riazunnisa et al., 2006), O_2_ generated by photosynthesis can be consumed by mitochondria without diffusing out of the cell (Lavergne, 1989) and functional evidence in Arabidopsis suggests H_2_O_2_ accumulation induces cooperation between plastids and mitochondria to alleviate PSI acceptor limitation (Fitzpatrick et al 2022). In the plant, increased light dependent respiratory CO_2_ evolution was seen when infiltrating Arabidopsis with MV. These experiments were consistent with a possible H_2_O_2_ priming of mitochondria to accept excess energy, potentially through malate shuttle. The MCFc mutations generated in this work affect both photosynthesis (Fig. 2) and respiration (Fig. 3). Thus, they alter the relationship between these two processes in mutant cells (Fig. 3A).

### Molecular players involved in organelle exchanges in diatoms

Despite functional studies (Bailleul et al., 2015), and modelling approaches (Levering et al. 2016, Broddrick et al. 2019) the molecular mechanisms governing plastid-mitochondria interactions in diatoms, and complex plastids in general, remain poorly understood. Previous research has identified bacteria-like plastidial nucleotide transporters (NTTs) in diatoms, which are localized in the plastid envelope (Ast et al. 2009, Gruber et Haferkamp 2019). These transporters may play a role in supplying nucleotides to the plastid for energy metabolism, but their exact energetic role is unclear, as they do not catalyze ATP/ADP exchange like in primary plastids (Ast et al., 2009; Chu et al., 2017).

Several other diatom transporters localized to the thylakoid or envelope membranes have been studied (reviewed in Marchand et al., 2018; Finazzi et al., 2015; Carretto et al., 2016). These transporters primarily include carriers of ions and a limited number of metabolites. However, their known activities and substrate specificities do not align with the functions investigated in this study.

Bioinformatic, molecular, and physiological analyses suggest that it is a member of the MCF (mitochondrial carrier family) group. Confocal imaging of GFP-tagged strains confirms that MCFc is specifically targeted to the plastid. While its precise substrate remains unidentified, the deletion of MCFc significantly impacts the accumulation of the key amino acid precursor 2-KG (α-ketoglutarate) and other amino acids. This disruption is likely due to impaired partitioning of aspartate, glutamate, asparagine, and arginine between the plastid and cytosol. Our analysis suggests different hypothesis to explain its function:

Hypothesis 1, (the most plausible hypothesis, Fig.4C) that would account for the different results described here would be that MCFc transports 2-KG inside the plastid (maybe against Glutamate). It would explain (i) a lower ETR (decreased electron sink in the plast), (ii), metabolomics data (see Results part), (iii) MIMS results (default in 2-KG import into the plastid would re-route this metabolite towards complete respiration in the TCA cycle) and (iv) it would be consistent with *in silico* predictions. In land plants, oxoglutarate/glutamate exchange is carried out by the combined action of DIT1 (2-KG_in_/malate_out_) and DIT2 (glutamate_out_/malate_in_) resulting in no net malate transport but net export of fixed nitrogen. The ortholog of DIT1 in *P. tricornutum* encoded by Phatr3_EG02645, shares domain homology with the Arabidopsis DIT1 2-oxoglutarate-malate exchanger. However, the gene model lacks an ER transit peptide signal and the diatom plastid targeting signal (Gruber et al., 2007). Additionally, the occurrence of plastid-localized enzymes in *P. tricornutum* that use malate as a substrate has not been found yet in the genome (Broddrick et al., 2019), raising doubts about a DIT1/DIT2 cooperation in diatoms.

Hypothesis 2 (Figure 4 C). The third putative substrate suggested by *in silico* analysis is citrate (Supplementary Figure 4, Supplementary Table 1). Citrate does not have any known documented metabolic role in the plastid in diatoms, nor in land plant. However, tricarboxylate transporters transport both citrate and isocitrate (Picault et al., 2002). A plastidial isocitrate dehydrogenase (catalyzing the reaction isocitrate + NADP+ <-> 2-KG + NADPH + CO_2_) has been identified in both land plants (see http://chlorokb.fr) and *P. tricornutum* (Huang et al., 2021). If this plastidial enzyme functions in the direction of isocitrate decarboxylation (Direction 1), then cytosolic isocitrate must be imported into the plastid. This pathway would provide an alternative source of 2-KG for nitrogen assimilation. The concomitant production of NADPH by isocitrate dehydrogenase — apparently unnecessary in the plastid during the light phase —may explain why the role of this enzyme is not yet fully understood. However, if the isocitrate dehydrogenase works in the opposite direction (Direction 2, 2-KG + NADPH + CO_2_ -> isocitrate + NADP^+^), it could serve as a mechanism to alleviate redox pressure in the plastid. Defective isocitrate import (isocitrate dehydrogenase operating in direction 1) could also explain MIMS results and metabolic profile. Regardless of the direction in which the enzyme operates, isocitrate transport across the plastid envelope is essential, although no such transporter has yet been identified.

In conclusion, this study underscores the intricate nature of intracellular metabolite transport and its connection to energy metabolism in diatoms. Although many key molecular players remain elusive, our integrated approach—encompassing molecular biology, bioinformatics, and physiology—provides valuable insights and lays a strong foundation for advancing our understanding of these unique organisms in the years to come.

## Author contributions

**Conceptualization:** Giovanni Finazzi.

**Coordination:** Giovanni Finazzi, Gilles Curien, Alisdair R. Fernie, Pierre Cardol, Denis Baurain, Eva-Mari Aro, and Dimitri Tolleter coordinated the research.

**Experiments:** Cécile Giustini, Davide Dal Bo, Mattia Storti, Mick Van Vlierberghe, Denis Baurain, Youjun Zhang, Duncan Fitzpatrick, Guillaume Allorent, Pascal Albanese, and Dimitri Tolleter performed the experiments.

**Manuscript Writing:** Giovanni Finazzi and Dimitri Tolleter drafted the manuscript with contributions from all authors.

**Review and Approval:** All authors reviewed and approved the final manuscript.

## Acknowledgments

We thank Richard Dorrell for kindly providing the MCFc::GFP mutant, Luigi and Ferdinando Palmieri for their helpful discussions and efforts on MCFc transporter assays and Chris Bowler for fruitful discussion in the initial phase of the project.

C.G., G.C., M.S., D.T. and G.F. acknowledge funding from the European Research Council ERC (Chloro-Mito; grant no. 833184). P.A. acknowledges funding from the European Union’s Horizon 2020 research and innovation program under the Marie Skłodowska-Curie grant agreement No 101066400 - PHOTO-LINK. G.F. acknowledge funds from the Plankton project (grant agreement 101099192). G.A. and E.M.A. acknowledge funding from the CNRS Momentum program and the Jane and Aatos Erkko Foundation, respectively.

## References

Alric J. 2010. Cyclic electron flow around photosystem I in unicellular green algae. Photosynth Res. 2010 Nov;106(1-2):47–56. doi: 10.1007/s11120-010-9566-4…

Altschul SF, Gish W, Miller W, Myers EW, Lipman DJ. 1990. Basic local alignment search tool. J Mol Biol. 1990 Oct 5;215(3):403–10. doi: 10.1016/S0022-2836(05)80360-2.

Asada K. 2000 The water-water cycle as alternative photon and electron sinks. Philos Trans R Soc Lond B Biol Sci. 2000 Oct 29;355(1402):1419–31. doi: 10.1098/rstb.2000.0703.

Ast, M., Gruber, A., Schmitz-Esser, S., Neuhaus, H. E., Kroth, P. G., Horn, M., & Haferkamp, I. (2009). Diatom plastids depend on nucleotide import from the cytosol. Proceedings of the National Academy of Sciences of the United States of America, 106(9), 3621–3626.

Bailleul B., Cardol P., Breyton C., Finazzi G. 2010. Electrochromism: a useful probe to study algal photosynthesis Photosynth. Res., 106, pp. 179–189

Bailleul B, Berne N, Murik O, et al. 2015. Energetic coupling between plastids and mitochondria drives CO2 assimilation in diatoms. Nature. 2015 Aug 20;524(7565):366–9. doi: 10.1038/nature14599…

Beckmann K, Messinger J, Badger MR, Wydrzynski T, Hillier W. 2009. On-line mass spectrometry: Membrane inlet sampling. Photosynth Res 102: 511–522

Berges, John A, Daniel J Franklin, and Paul J Harrison. 2001. Evolution Of An Artificial Seawater Medium : Improvements In Enriched Seawater, Artificial Water Over The Last Two Decades. 1145:1138–45.

Bilger W., Björkman O., 1990. Role of the xanthophyll cycle in photoprotection elucidated by measurements of light-induced absorbance changes, fluorescence and photosynthesis in leaves of Hedera canariensis. Photosynth. Res. 25, 173–185.

Broddrick JT, D. N, Smith SR, et al. 2019. Cross-compartment metabolic coupling enables flexible photoprotective mechanisms in the diatom Phaeodactylum tricornutum. New Phytol. 2019 May;222(3):1364–1379. doi: 10.1111/nph.15685…

Burlacot A., Burlacot F., Li-Beisson Y., Peltier G. 2020. Membrane Inlet Mass Spectrometry : A powerful tool for algal research. Frontiers in Plant Science

Busch FA, Sage TL, Cousins AB, Sage RF. 2013. C3 plants enhance rates of photosynthesis by reassimilating photorespired and respired CO2. Plant Cell Environ. 2013 Jan;36(1):200–12. doi: 10.1111/j.1365-3040.2012.02567.x…

Cardol P, Alric J, Girard-Bascou J, Franck F, Wollman FA, Finazzi G. 2009. Impaired respiration discloses the physiological significance of state transitions in Chlamydomonas. Proc Natl Acad Sci U S A. 2009 Sep 15;106(37):15979–84. doi: 10.1073/pnas.0908111106. Erratum in: Proc Natl Acad Sci U S A. 2019 Apr 2;116(14):7150. doi: 10.1073/pnas.1903574116.

Carraretto L, Teardo E, Checchetto V, Finazzi G, Uozumi N, Szabo I. 2016. Ion Channels in Plant Bioenergetic Organelles, Chloroplasts and Mitochondria: From Molecular Identification to Function. Molecular Plant 9-3 pp371–395 10.1016/j.molp.2015.12.004.

Chu, L., Gruber, A., Ast, M., et al. 2017. Shuttling of (deoxy-) purine nucleotides between compartments of the diatom Phaeodactylum tricornutum. The New phytologist, 213(1), 193–205.

Curien G, Flori S, Villanova V, et al. 2016. The Water to Water Cycles in Microalgae. Plant Cell Physiol. 2016 Jul;57(7):1354–1363. doi: 10.1093/pcp/pcw048…

Ellis, R. J. (2010). Tackling unintelligent design. Nature, 463(7278), 164–165.

Felsenstein J. 1978. Cases in which parsimony or compatibility methods will be positively misleading. Syst Zool 27: 401–410.

Finazzi G, Petroutsos D, Tomizioli M, Flori S, Sautron E, Villanova V, Rolland N, Seigneurin-Berny D. 2015. Ions channels/transporters and chloroplast regulation. Cell Calcium. 2015 Jul;58(1):86–97. doi: 10.1016/j.ceca.2014.10.002.

Fitzpatrick Duncan, Aro Eva-Mari, Tiwari Arjun, True oxygen reduction capacity during photosynthetic electron transfer in thylakoids and intact leaves, Plant Physiology, Volume 189, Issue 1, May 2022, Pages 112–128,

Giustini, C., Angulo, J., Courtois, F., Allorent, G. 2024. Targeted Gene Editing of Nuclear-Encoded Plastid Proteins in Phaeodactylum tricornutum via CRISPR/Cas9. In: Maréchal, E. (eds) Plastids. Methods in Molecular Biology, vol 2776. Humana, New York, NY.

Gouy R, Baurain D, Philippe H. 2015. Rooting the tree of life: the phylogenetic jury is still out. Philos Trans R Soc Lond B Biol Sci 370: 20140329.

Gruber A., Rocap G., Kroth P.G., Armbrust E.V., Mock T. 2015. Plastid proteome prediction for diatoms and other algae with secondary plastids of the red lineage”. In: The Plant Journal 81.3, pp. 519–528. doi: 10.1111/tpj.12734. url: https://doi.org/10.1111/tpj.12734.

Gruber A, Haferkamp I. Nucleotide Transport and Metabolism in Diatoms. Biomolecules. 2019 Nov 21;9(12):761. doi: 10.3390/biom9120761.

Harrison, Paul J, Rosemary E Waters, and F J R Taylor. 1980. A Broad Spectrum Artificial Sea Water Medium For Coastal And Open Ocean Phytoplankton. Journal of Phycology 16(1): 28–35.130.

Hoefnagel M.H.N., Atkin O.K., et Wiskich J.T. 1998. Interdependence between chloroplasts and mitochondria in the light and the dark. Biochimica et Biophysica Acta (BBA)-Bioenergetics, vol. 1366, no 3, p. 235–255.

Irisarri I, Baurain D, Brinkmann H, Delsuc F, Sire JY, Kupfer A, Petersen J, Jarek M, Meyer A, Vences M et al. 2017. Phylotranscriptomic consolidation of the jawed vertebrate timetree. Nat Ecol Evol 1: 1370–1378.

Joliot, P., & Joliot, A. (1989). Characterization of linear and quadratic electrochromic probes in Chlorella sorokiniana and Chlamydomonas reinhardtii. Biochimica et Biophysica Acta (BBA)-Bioenergetics, 975(3), 355–360.

M.J. Johnson, A.V. Ruban (2014) Rethinking the existence of a steady-state Δψ component of the proton motive force across plant thylakoid membranes Photosynth Res 119 pp :233–242

Katoh K, Standley DM. 2013. MAFFT multiple sequence alignment software version 7: improvements in performance and usability. Mol Biol Evol 30:772–780.

Kempen, Michel van, Stephanie S. Kim, Charlotte Tumescheit, Milot Mirdita, Jeongjae Lee, Cameron L. M. Gilchrist, Johannes Söding, and Martin Steinegger. 2024. “Fast and Accurate Protein Structure Search with Foldseek.” Nature Biotechnology 42 (2): 243–46. 10.1038/s41587-023-01773-0.

Kinoshita H, Nagasaki J, Yoshikawa A. et al. 2011. The Chloroplastic 2-Oxoglutarate/Malate Transporter Has Dual Function as the Malate Valve and in Carbon/Nitrogen Metabolism. The Plant Journal 65(1): 15–26.

Hoefnagel, Marcel H.N, Owen K Atkin, and Joseph T Wiskich. 1998. “Interdependence between Chloroplasts and Mitochondria in the Light and the Dark.” Biochimica et Biophysica Acta (BBA) - Bioenergetics 1366 (3): 235–55. 10.1016/s0005-2728(98)00126-1

Hoguin, A., Yang, F., Groisillier, A. et al. 2023. The model diatom Phaeodactylum tricornutum provides insights into the diversity and function of microeukaryotic DNA methyltransferases. Commun Biol 6, 253

Huang, Shiping, Jiaxin Zhao, Wenjing Li, Peng Wang, Zhenglian Xue, and Guoping Zhu. 2021. “Biochemical and Phylogenetic Characterization of a Novel NADP+-Specific Isocitrate Dehydrogenase From the Marine Microalga Phaeodactylum Tricornutum.” Frontiers in Molecular Biosciences 8:702083.

Huppe HC, and Turpin DH, Integration of carbon and nitrogen metabolism in plant and algae cells, Annual review of Plant Biology 45:577–607 (1994) 10.1146/annurev.pp.45.060194.003045

Keeling PJ, Burki F, Wilcox HM et al. 2014. The Marine Microbial Eukaryote Transcriptome Sequencing Project (MMETSP): illuminating the functional diversity of eukaryotic life in the oceans through transcriptome sequencing. PLoS Biol. 2014;12(6):e1001889.

Lavergne J. 1989. Mitochondrial responses to intracellular pulses of photosynthetic oxygen. Proc Natl Acad Sci U S A. 1989 Nov;86(22):8768–72. doi: 10.1073/pnas.86.22.8768.

Le QS, Gascuel O, Lartillot N. 2008. Empirical profile mixture models for phylogenetic reconstruction. Bioinformatics 24: 2317–2323.

Le SQ, Dang CC, Gascuel O. 2012. Modeling protein evolution with several amino acid replacement matrices depending on site rates. Mol Biol Evol 29: 2921–2936.

Lemaire C, Wollman FA, Bennoun P. Restoration of phototrophic growth in a mutant of Chlamydomonas reinhardtii in which the chloroplast atpB gene of the ATP synthase has a deletion: an example of mitochondria-dependent photosynthesis. Proc Natl Acad Sci U S A. 1988 Mar;85(5):1344–8. doi: 10.1073/pnas.85.5.1344.

Levering J, Broddrick J, Dupont CL, Peers G, Beeri K, Mayers J, Gallina AA, Allen AE, Palsson BO, Zengler K. 2016. Genome-Scale Model Reveals Metabolic Basis of Biomass Partitioning in a Model Diatom. PLoS One. 2016 May 6;11(5):e0155038. doi: 10.1371/journal.pone.0155038.

Letunic I and Bork P (2024) Interactive Tree of Life (iTOL) v6: recent updates to the phylogenetic tree display and annotation tool. Nucleic Acids Res doi: 10.1093/nar/gkae268

Li L, Wang S, Wang H, Sahu SK, Marin B, Li H, Xu Y, Liang H, Li Z, Cheng S et al. 2020. The genome of Prasinoderma coloniale unveils the existence of a third phylum within green plants. Nat Ecol Evol 4: 1220–1231.

Liu S., Yang SM, Bowler C., Obornik M., Dorrell R., Dynamic relocalization and divergent expression of a major facilitator carrier subfamily in diatoms. 2024 bioRxiv 10.1101/2024.12.16.628690

Marchand J, Heydarizadeh P, Schoefs B, Spetea C. Ion and metabolite transport in the chloroplast of algae: lessons from land plants. Cell Mol Life Sci. 2018 Jun;75(12):2153–2176. doi: 10.1007/s00018-018-2793-0. PMID:29541792;.

Mehler AH. 1951. Studies on reactions of illuminated chloroplasts. I. Mechanism of the reduction of oxygen and other Hill reagents. Arch Biochem Biophys. 1951 Aug;33(1):65–77. doi: 10.1016/0003-9861(51)90082-3.

Merchant SS, Prochnik SE, Vallon O, et al. 2007. The Chlamydomonas Genome Reveals the Evolution of Key Animal and Plant Functions. Science 318, 245–250 (2007). DOI:.

Minh BQ, Schmidt HA, Chernomor O, Schrempf D, Woodhams MD, von Haeseler A, Lanfear R. 2020. IQ-TREE 2: New Models and Efficient Methods for Phylogenetic Inference in the Genomic Era. Mol Biol Evol 37: 1530–1534.

Murik O, Tirichine L, Prihoda J, Thomas Y, Araújo WL, Allen AE, Fernie AR, Bowler C. 2019. Downregulation of mitochondrial alternative oxidase affects chloroplast function, redox status and stress response in a marine diatom. New Phytol. 2019 Feb;221(3):1303–1316. doi: 10.1111/nph.15479…

Obata, T., Schoenefeld, S., Krahnert, I., et al. 2013. Gas-chromatography mass-spectrometry (GC-MS) based metabolite profiling reveals mannitol as a major storage carbohydrate in the coccolithophorid alga Emiliania huxleyi. Metabolites, 3(1), 168–184.

Ort DR, Baker NR. 2002. A photoprotective role for O(2) as an alternative electron sink in photosynthesis? Curr Opin Plant Biol. 2002 Jun;5(3):193–8. doi: 10.1016/s1369-5266(02)00259-5.

Palmieri, Ferdinando, and Ciro Leonardo Pierri. 2010. “Structure and Function of Mitochondrial Carriers – Role of the Transmembrane Helix P and G Residues in the Gating and Transport Mechanism.” FEBS Letters 584 (9): 1931–39. 10.1016/j.febslet.2009.10.063.

Palmieri F, Pierri CL, De Grassi A, Nunes-Nesi A, Fernie AR. 2011. Evolution, structure and function of mitochondrial carriers: a review with new insights. Plant J. 2011 Apr;66(1):161–81. doi: 10.1111/j.1365-313X.2011.04516.x.

Palmieri F. 2013. The mitochondrial transporter family SLC25: identification, properties and physiopathology. Mol Aspects Med. 2013 Apr-Jun;34(2-3):465–84. doi: 10.1016/j.mam.2012.05.005…

Petersen J, Förster K, Turina P, Gräber P. 2012. Comparison of the H+/ATP ratios of the H+-ATP synthases from yeast and from chloroplast. Proc Natl Acad Sci U S A. 2012 Jul 10;109(28):11150–5. doi: 10.1073/pnas.1202799109…

Petersen J, Ludewig AK, Michael V, Bunk B, Jarek M, Baurain D, Brinkmann H. 2014. Chromera velia, endosymbioses and the rhodoplex hypothesis--plastid evolution in cryptophytes, alveolates, stramenopiles, and haptophytes (CASH lineages). Genome Biol Evol 6: 666–684.

Petersen TN, Brunak S, von Heijne G, Nielsen H. 2011. SignalP 4.0: discriminating signal peptides from transmembrane regions. Nat Methods. 2011 Sep 29;8(10):785–6. doi: 10.1038/nmeth.1701.

Picault N., Palmieri L., Pisano I., et al., 2002. Identification of a novel transporter for dicarboxylates and tricarboxylates in plant mitochondria. Bacterial expression, reconstitution, functional characterization, and tissue distribution, J Biol Chem Jul 5;277(27):24204–11. Doi: 10.1074/jbc.M202702200

Prihoda J., Tanaka A., de Paula Wilson B. M., et al. 2012, Chloroplast-mitochondria cross-talk in diatoms, Journal of Experimental Botany, Volume 63, Issue 4, February 2012, Pages 1543–1557,

Achal R., Murik O., Bowler C., and Tirichine L. 2016. PhytoCRISP-Ex: A Web-Based and Stand-Alone Application to Find Specific Target Sequences for CRISPR/CAS Editing. BMC bioinformatics 17(1): 261. https://www.ncbi.nlm.nih.gov/pubmed/27363443.

Riazunnisa, K., Padmavathi, L., Bauwe, H., & Raghavendra, A. S. 2006. Markedly low requirement of added CO2 for photosynthesis by mesophyll protoplasts of pea (Pisum sativum): possible roles of photorespiratory CO2 and carbonic anhydrase. Physiologia Plantarum, 128(4), 763–772.

Ruprecht, Jonathan J., and Edmund R.S. Kunji. 2020. The SLC25 Mitochondrial Carrier Family: Structure and Mechanism. Trends in Biochemical Sciences 45 (3): 244–58. 10.1016/j.tibs.2019.11.001.

Satre M, Mattei S, Aubry L, Gaudet P, Pelosi L, Brandolin G, Klein G. 2007. Mitochondrial carrier family: repertoire and peculiarities of the cellular slime mould Dictyostelium discoideum. Biochimie. 2007 Sep;89(9):1058–69. doi: 10.1016/j.biochi.2007.03.004…

Seydoux, C., Storti, M., Giovagnetti, V., et al. 2022. Impaired photoprotection in Phaeodactylum tricornutum KEA3 mutants reveals the proton regulatory circuit of diatoms light acclimation. The New phytologist, 234(2), 578–591. 10.1111/nph.18003

Shikanai T. Cyclic electron transport around photosystem I: genetic approaches. Annu Rev Plant Biol. 2007;58:199–217. doi: 10.1146/annurev.arplant.58.091406.110525.

Simion P, Philippe H, Baurain D, et al. 2017. A Large and Consistent Phylogenomic Dataset Supports Sponges as the Sister Group to All Other Animals. Curr Biol 27: 958–967.

Smith, S.R., Dupont, C.L., McCarthy, J.K. et al. 2019. Evolution and regulation of nitrogen flux through compartmentalized metabolic networks in a marine diatom. Nat Commun 10, 4552 (2019). 10.1038/s41467-019-12407-y

Varadi, Mihaly, Damian Bertoni, Paulyna Magana, Urmila Paramval, Ivanna Pidruchna, Malarvizhi Radhakrishnan, Maxim Tsenkov, et al. 2024. “AlphaFold Protein Structure Database in 2024: Providing Structure Coverage for over 214 Million Protein Sequences.” Nucleic Acids Research 52 (D1): D368–75.

Van Vlierberghe M, Philippe H, Baurain D. 2021a. Broadly sampled orthologous groups of eukaryotic proteins for the phylogenetic study of plastid-bearing lineages. BMC Research Notes 14.

Van Vlierberghe M, Di Franco A, Philippe H, Baurain D. 2021b. Decontamination, pooling and dereplication of the 678 samples of the Marine Microbial Eukaryote Transcriptome Sequencing Project. BMC Research Notes 14.

Villanova, V., Fortunato, A.E., Singh, D., et al. 2017. Investigating mixotrophic metabolism in the model diatom Phaeodactylum tricornutum. Philosophical Transactions of the Royal Society B: Biological Sciences, 372(1728), p. 20160404.

Villanova V, Singh D, Pagliardini J, et al. 2021 Boosting Biomass Quantity and Quality by Improved Mixotrophic Culture of the Diatom Phaeodactylum tricornutum. Front Plant Sci. 2021 Apr 9;12:642199. doi: 10.3389/fpls.2021.642199. eCollection 2021.

Uwizeye, Clarisse, Johan Decelle, Pierre-Henri Jouneau, Serena Flori, Benoit Gallet, Jean-Baptiste Keck, Davide Dal Bo, et al. 2021. “Morphological Bases of Phytoplankton Energy Management and Physiological Responses Unveiled by 3D Subcellular Imaging.” Nature Communications 12 (1). 10.1038/s41467-021-21314-0.

Zehr JP, Kudela RM. Ocean science. Photosynthesis in the open ocean. Science. 2009 Nov 13;326(5955):945–6. doi: 10.1126/science.1181277.

