## Supplementary Figure 1A for "A Mitochondrially Derived Plastidial Transporter Regulates Photosynthesis in the Diatom *Phaeodactylum tricornutum*"

A

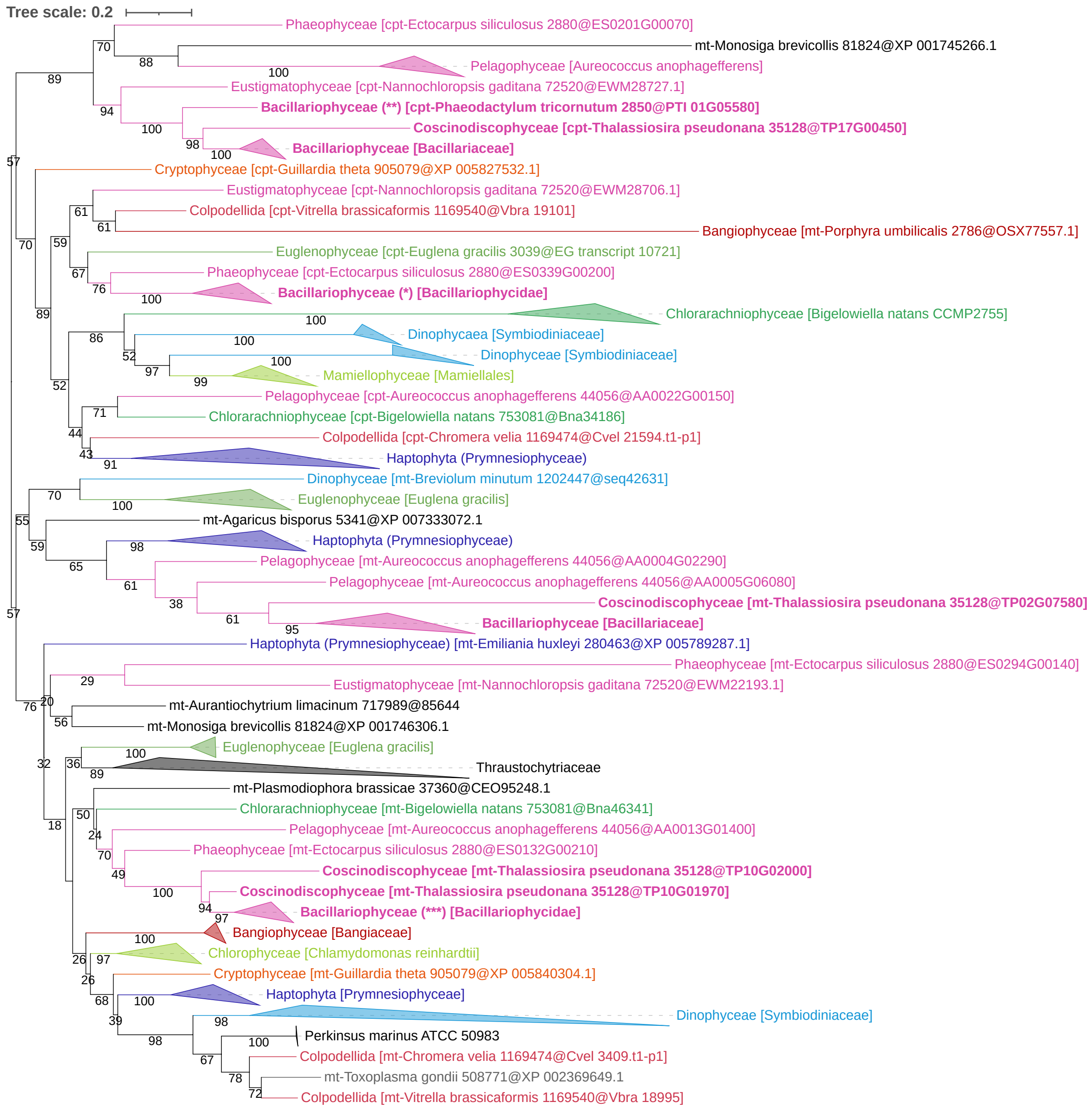
Tree scale: 0.2
Phaeophyceae [cpt-Ectocarpus siliculosus 2880@ES0201G00070]
mt-Monosiga brevicollis 81824@XP 001745266.1
Pelagophyceae [Aureococcus anophagefferens]
Eustigmatophyceae [cpt-Nannochloropsis gaditana 72520@EWM28727.1]
Bacillariophyceae (**) [cpt-Phaeodactylum tricornutum 2850@PTI 01G05580]
Coscinodiscophyceae [cpt-Thalassiosira pseudonana 35128@TP17G00450]
Bacillariophyceae [Bacillariaceae]
Cryptophyceae [cpt-Guillardia theta 905079@XP 005827532.1]
Eustigmatophyceae [cpt-Nannochloropsis gaditana 72520@EWM28706.1]
Colpodellida [cpt-Vitrella brassicaformis 1169540@Vbra 19101]
Bangiophyceae [mt-Porphyra umbilicalis 2786@OSX77557.1]
Euglenophyceae [cpt-Euglena gracilis 3039@EG transcript 10721]
Phaeophyceae [cpt-Ectocarpus siliculosus 2880@ES0339G00200]
Bacillariophyceae (*) [Bacillariophycidae]
Chlorarachniophyceae [Bigelowiella natans CCMP2755]
Dinophycaea [Symbiodiniaceae]
Dinophyceae [Symbiodiniaceae]
Mamiellophyceae [Mamiellales]
Pelagophyceae [cpt-Aureococcus anophagefferens 44056@AA0022G00150]
Chlorarachniophyceae [cpt-Bigelowiella natans 753081@Bna34186]
Colpodellida [cpt-Chromera velia 1169474@Cvel 21594.t1-p1]
Haptophyta (Prymnesiophyceae)
Dinophyceae [mt-Breviolum minutum 1202447@seq42631]
Euglenophyceae [Euglena gracilis]
mt-Agaricus bisporus 5341@XP 007333072.1
Haptophyta (Prymnesiophyceae)
Pelagophyceae [mt-Aureococcus anophagefferens 44056@AA0004G02290]
Pelagophyceae [mt-Aureococcus anophagefferens 44056@AA0005G06080]
Coscinodiscophyceae [mt-Thalassiosira pseudonana 35128@TP02G07580]
Bacillariophyceae [Bacillariaceae]
Haptophyta (Prymnesiophyceae) [mt-Emiliania huxleyi 280463@XP 005789287.1]
Phaeophyceae [mt-Ectocarpus siliculosus 2880@ES0294G00140]
Eustigmatophyceae [mt-Nannochloropsis gaditana 72520@EWM22193.1]
mt-Aurantiochytrium limacinum 717989@85644
mt-Monosiga brevicollis 81824@XP 001746306.1
Euglenophyceae [Euglena gracilis]
Thraustochytriaceae
mt-Plasmodiophora brassicae 37360@CEO95248.1
Chlorarachniophyceae [mt-Bigelowiella natans 753081@Bna46341]
Pelagophyceae [mt-Aureococcus anophagefferens 44056@AA0013G01400]
Phaeophyceae [mt-Ectocarpus siliculosus 2880@ES0132G00210]
Coscinodiscophyceae [mt-Thalassiosira pseudonana 35128@TP10G02000]
Coscinodiscophyceae [mt-Thalassiosira pseudonana 35128@TP10G01970]
Bacillariophyceae (***) [Bacillariophycidae]
Bangiophyceae [Bangiaceae]
Chlorophyceae [Chlamydomonas reinhardtii]
Cryptophyceae [mt-Guillardia theta 905079@XP 005840304.1]
Haptophyta [Prymnesiophyceae]
Dinophyceae [Symbiodiniaceae]
Perkinsus marinus ATCC 50983
Colpodellida [mt-Chromera velia 1169474@Cvel 3409.t1-p1]
mt-Toxoplasma gondii 508771@XP 002369649.1
Colpodellida [mt-Vitrella brassicaformis 1169540@Vbra 18995]

1

**B**

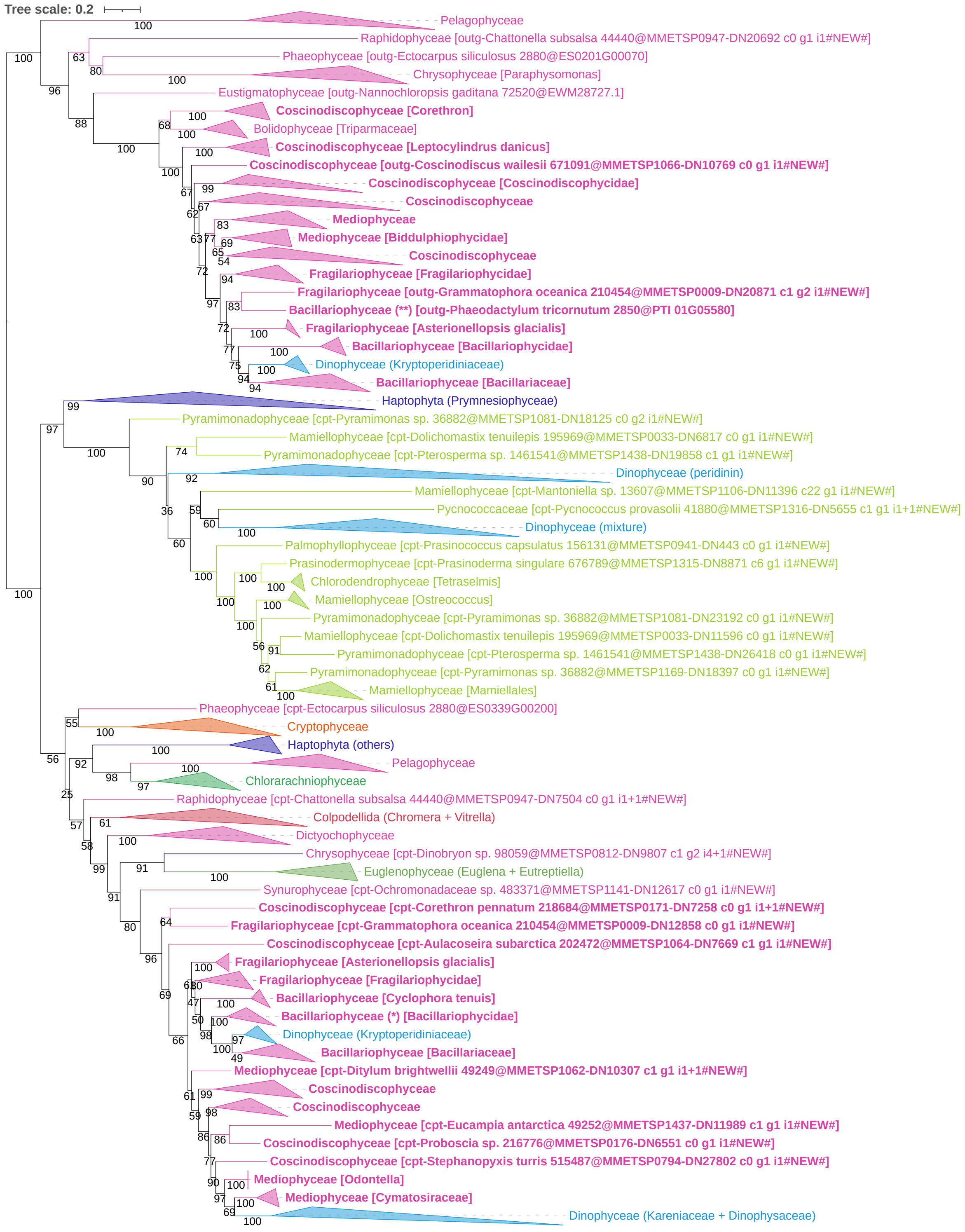
Tree scale: 0.2
Pelagophyceae
Raphidophyceae [outg-Chattonella subsalsa 44440@MMETSP0947-DN20692 c0 g1 i1#NEW#]
Phaeophyceae [outg-Ectocarpus siliculosus 2880@ES0201G00070]
Chrysophyceae [Paraphysomonas]
Eustigmatophyceae [outg-Nannochloropsis gaditana 72520@EWM28727.1]
Coscinodiscophyceae [Corethron]
Bolidophyceae [Triparmaceae]
Coscinodiscophyceae [Leptocylindrus danicus]
Coscinodiscophyceae [outg-Coscinodiscus wailesii 671091@MMETSP1066-DN10769 c0 g1 i1#NEW#]
Coscinodiscophyceae [Coscinodiscophycidae]
Coscinodiscophyceae
Mediophyceae
Mediophyceae [Biddulphiophycidae]
Coscinodiscophyceae
Fragilariophyceae [Fragilariophycidae]
Fragilariophyceae [outg-Grammatophora oceanica 210454@MMETSP0009-DN20871 c1 g2 i1#NEW#]
Bacillariophyceae (**) [outg-Phaeodactylum tricornutum 2850@PTI 01G05580]
Fragilariophyceae [Asterionellopsis glacialis]
Bacillariophyceae [Bacillariophycidae]
Dinophyceae (Kryptoperidiniaceae)
Bacillariophyceae [Bacillariaceae]
Haptophyta (Prymnesiophyceae)
Pyramimonadophyceae [cpt-Pyramimonas sp. 36882@MMETSP1081-DN18125 c0 g2 i1#NEW#]
Mamiellophyceae [cpt-Dolichomastix tenuilepis 195969@MMETSP0033-DN6817 c0 g1 i1#NEW#]
Pyramimonadophyceae [cpt-Pterosperma sp. 1461541@MMETSP1438-DN19858 c1 g1 i1#NEW#]
Dinophyceae (peridinin)
Mamiellophyceae [cpt-Mantoniella sp. 13607@MMETSP1106-DN11396 c22 g1 i1#NEW#]
Pycnococcaceae [cpt-Pycnococcus provasolii 41880@MMETSP1316-DN5655 c1 g1 i1+1#NEW#]
Dinophyceae (mixture)
Palmophyllophyceae [cpt-Prasinococcus capsulatus 156131@MMETSP0941-DN443 c0 g1 i1#NEW#]
Prasinodermophyceae [cpt-Prasinoderma singulare 676789@MMETSP1315-DN8871 c6 g1 i1#NEW#]
Chlorodendrophyceae [Tetraselmis]
Mamiellophyceae [Ostreococcus]
Pyramimonadophyceae [cpt-Pyramimonas sp. 36882@MMETSP1081-DN23192 c0 g1 i1#NEW#]
Mamiellophyceae [cpt-Dolichomastix tenuilepis 195969@MMETSP0033-DN11596 c0 g1 i1#NEW#]
Pyramimonadophyceae [cpt-Pterosperma sp. 1461541@MMETSP1438-DN26418 c0 g1 i1#NEW#]
Pyramimonadophyceae [cpt-Pyramimonas sp. 36882@MMETSP1169-DN18397 c0 g1 i1#NEW#]
Mamiellophyceae [Mamiellales]
Phaeophyceae [cpt-Ectocarpus siliculosus 2880@ES0339G00200]
Cryptophyceae
Haptophyta (others)
Pelagophyceae
Chlorarachniophyceae
Raphidophyceae [cpt-Chattonella subsalsa 44440@MMETSP0947-DN7504 c0 g1 i1+1#NEW#]
Colpodellida (Chromera + Vitrella)
Dictyochophyceae
Chrysophyceae [cpt-Dinobryon sp. 98059@MMETSP0812-DN9807 c1 g2 i4+1#NEW#]
Euglenophyceae (Euglena + Eutreptiella)
Synurophyceae [cpt-Ochromonadaceae sp. 483371@MMETSP1141-DN12617 c0 g1 i1#NEW#]
Coscinodiscophyceae [cpt-Corethron pennatum 218684@MMETSP0171-DN7258 c0 g1 i1+1#NEW#]
Fragilariophyceae [cpt-Grammatophora oceanica 210454@MMETSP0009-DN12858 c0 g1 i1#NEW#]
Coscinodiscophyceae [cpt-Aulacoseira subarctica 202472@MMETSP1064-DN7669 c1 g1 i1#NEW#]
Fragilariophyceae [Asterionellopsis glacialis]
Fragilariophyceae [Fragilariophycidae]
Bacillariophyceae [Cyclophora tenuis]
Bacillariophyceae (*) [Bacillariophycidae]
Dinophyceae (Kryptoperidiniaceae)
Bacillariophyceae [Bacillariaceae]
Mediophyceae [cpt-Ditylum brightwellii 49249@MMETSP1062-DN10307 c1 g1 i1+1#NEW#]
Coscinodiscophyceae
Coscinodiscophyceae
Mediophyceae [cpt-Eucampia antarctica 49252@MMETSP1437-DN11989 c1 g1 i1#NEW#]
Coscinodiscophyceae [cpt-Proboscia sp. 216776@MMETSP0176-DN6551 c0 g1 i1#NEW#]
Coscinodiscophyceae [cpt-Stephanopyxis turris 515487@MMETSP0794-DN27802 c0 g1 i1#NEW#]
Mediophyceae [Odontella]
Mediophyceae [Cymatosiraceae]
Dinophyceae (Kareniaceae + Dinophysaceae)

1

**C**

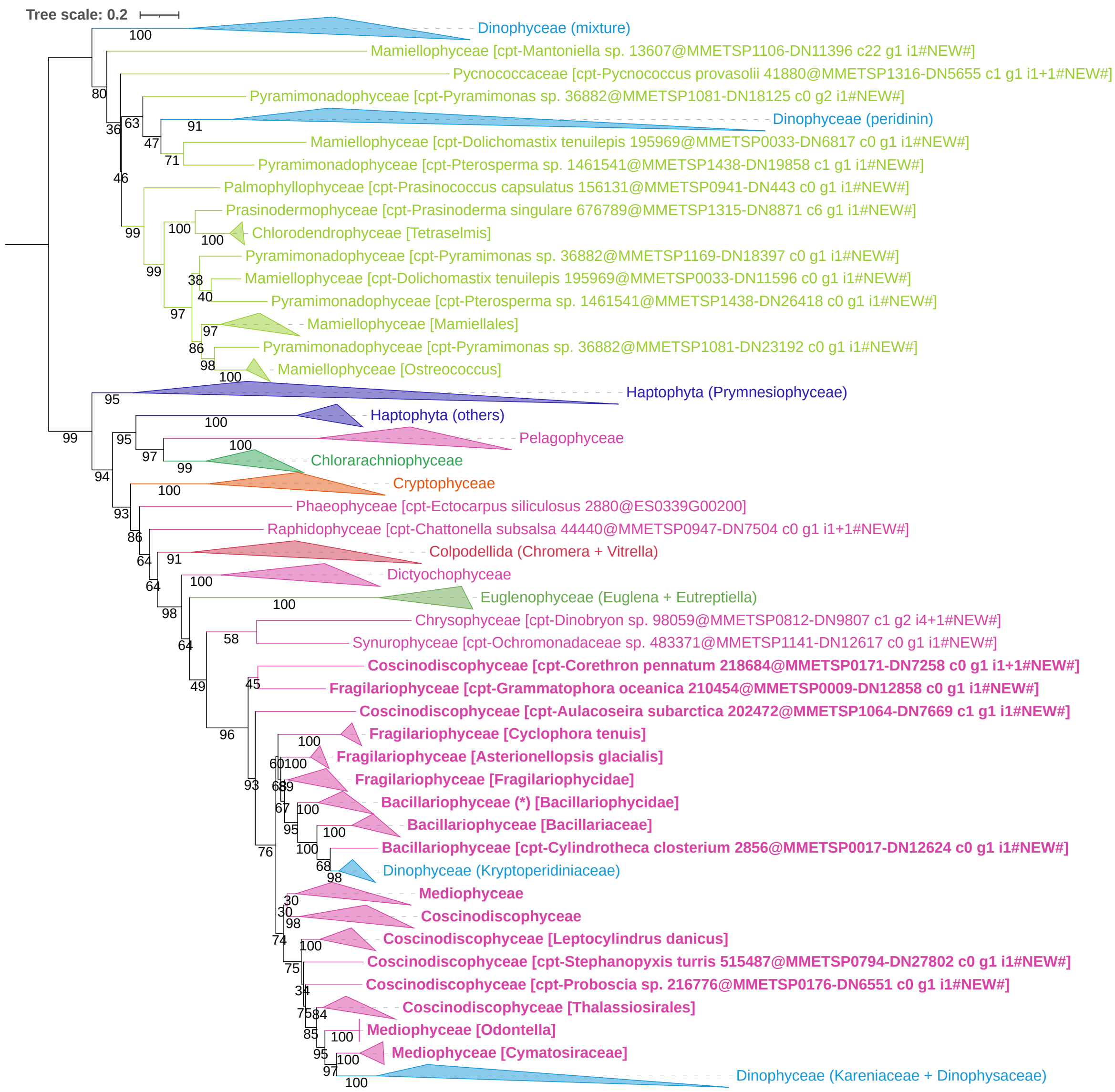


**Supplementary Figure 1.** Phylogenetic analyses of the MCFc-containing orthogroups (from Van Vlierberghe et al. 2021a). All trees were inferred under the C20 model. Monophyletic groups are collapsed at the family level and colored by taxonomic affiliation (light green: green algae, darker greens: chlorarachniophytes and euglenids, orange: cryptophytes, violet: haptophytes, pink: ochrophytes, blue: dinophytes, red: colpodellids). Diatoms are shown in boldface and the three groups each including one *Phaeodactylum* sequence (Phatr3_J46742, Phatr3_J42874 and Phatr3_J46612) are denoted by one, two and three asterisks (*), respectively. Numerical values at nodes are ultrafast bootstrap support (UFBS). Tree scales are in amino-acid substitutions per site. **(A)** Tree of the complete yet non-enriched orthogroup (OG0001460) featuring all three gene copies (92 sequences). One sequence of *Porphyra umbilicalis* is present but has a very long branch, and BLAST searches revealed a closer relationship to the other two red algal sequences present in the mitochondrial copy of the tree, suggesting a phylogenetic artifact. Along with the sequence of *Porphyra*, another sequence (of the choanoflagellate *Monosiga brevicolis*) shows a very long branch within the ochrophyte-specific copy, indicating that both sequences needed to be removed to avoid long-branch attraction (LBA) artifacts (Felsenstein 1978, Gouy et al. 2015). **(B)** Tree of the two PS subtrees (OG0001460-8) after transcriptomic enrichment and with the ochrophyte-specific copy used as the outgroup (286 sequences), from which two newly added sequences showing very long branches had to be removed too (the ochrophyte *Bolidomonas pacifica* and the haptophyte *Pavlova gyrans*). However, in such a scenario, green algae would be identified as receivers of a red gene, which here is implausible because the former are monophyletic (except for LGTs into dinophytes) and solely represented by a diverse array of basal lineages (Prasinodermophyta [Li et al. 2020)) and "prasinophytes", of which several display two or more distinct MCFc gene copies). **(C)** Tree fousing on the MCFc gene copy after pruning the ochrophyte-specific copy and rooted on the green algae (201 sequences). The latter tree is the same as the one shown in Fig. 1b, but with more complete labels and arranged in a rectangular view. Contrary to Fig, S1B, it still includes *Pavlova gyrans* (misplaced but hidden in the "Prymnesiophyceae" group).

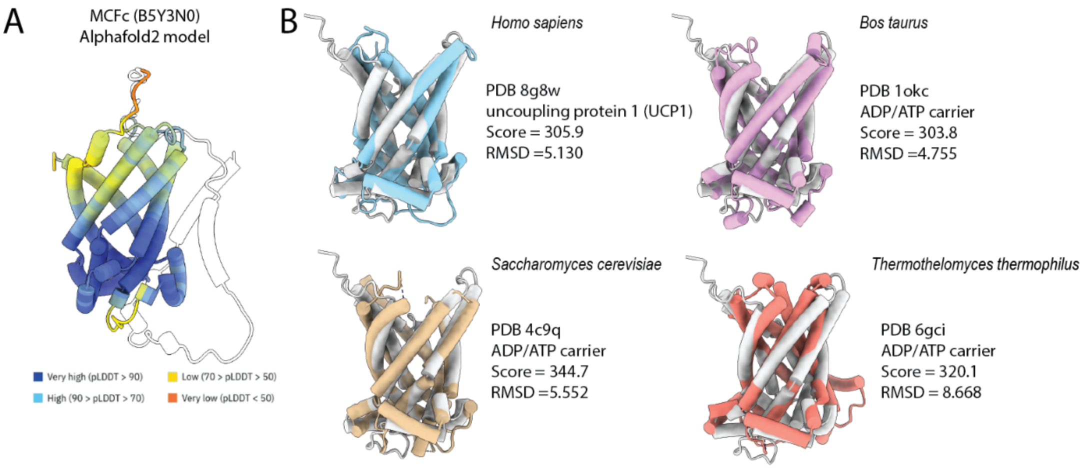


**Supplementary Figure 2**. Alphafold2 model of MCFc (A) with the putative signal peptide transparenct and redidues color according to the pLDDT confidence score. Structural alignment of MCFc predicted structure with known MCF mitochondrial carriers in the PDB repository. Fot each is indicated the specie, the PDB accession, the annotated function, the cumulative Root mean square deviation of atomic positions in Angstroms (RMSD) for the whole sequence and the overall alignment score. The score is calculated as: 0.70(residue similarity score, BLOSUM-62 matrix) + 0.30(secondary structure score) – gap penalties. Figures and alignments were performed in ChimeraX

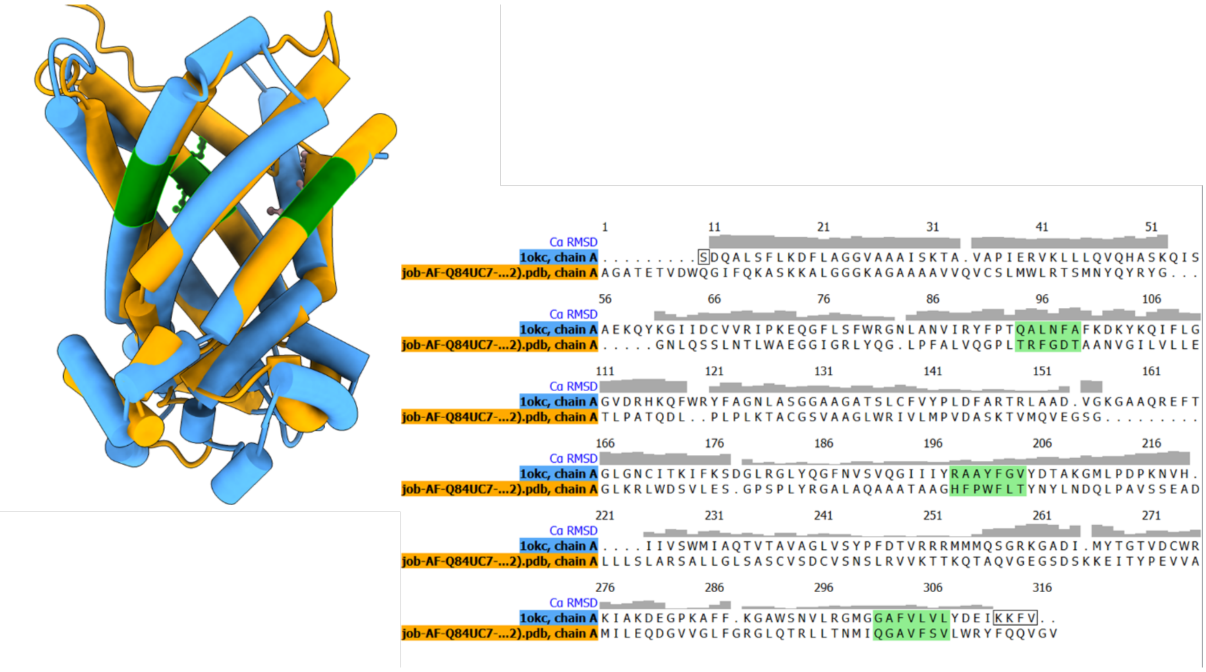


**Supplementary Figure 3**. Structural alignment, based on sequence proximity in the two compared structures used in supplementary Table 1. Motifs that are important for substrate selectivity in the cytoplasmic network are highlighted in green both in on the structure and on the pairwise sequence alignment. The Root mean square deviation of atomic positions in Angstroms (RMSD) for each amino acid backbone αCarbon is indicated. Figures and alignements were performed in ChimeraX (UCSF ChimeraX: Tools for structure building and analysis. Meng EC, Goddard TD, Pettersen EF, Couch GS, Pearson ZJ, Morris JH, Ferrin TE. Protein Sci. 2023 Nov;32(11):e4792.)

| | TM1-3-5 | TM2-4-6 | TOTAL |
|---|---|---|---|
| ***AGC, aspartate glutamate carrier*** | 13 | 5 | 18 |
| ***APC, ATP-Mg/Pi carrier*** | 14 | 3 | 17 |
| ***CIC, citrate (tricarboxylate) carrier*** | 13 | 3 | 16 |
| ***AAC, ADP/ATP carrier*** | 10 | 5 | 15 |
| ***CAC, carnitine-acylcarnitine carrier*** | 12 | 2 | 14 |
| ***OGC, oxoglutarate carrier*** | 11 | 1 | 12 |
| ***TPC, thiamine pyrophosphate carrier*** | 8 | 4 | 12 |
| ***PIC, phosphate carrier*** | 6 | 2 | 8 |
| ***UCP1, uncoupling protein.*** | 5 | 1 | 6 |

**Supplementary Table 1** . Table scoring the motif similarity in the six Trans membrane (TM) helixes (+2 if AA is same in the same position, +1 if flanking) according the alignment in supplementary figure 3. Each structure of the different transporters here listed was aligned and scored relative to the putative MCFc structure.

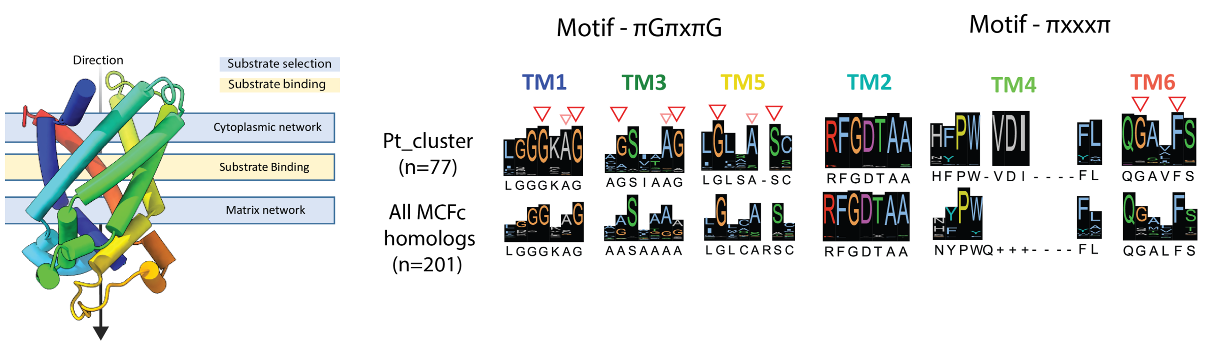


**Supplementary Figure 4**. Motif analysis of key position for substrate specificity in MCFc based on the alignment used for the phylogenetic tree in Figure 1B. MCFc cluster contains 77 sequences (includes Pt_MCFc) and for comparison the whole set of sequences (201 sequences, Supplementary Figure 1C). Arrows marks conserved sites matching for aspartate/glutamate, ATP-Mg/Pi or citrate carriers, if applicable. When no arrows are indicated it means that no similarity was found. ‘π’ is the one letter code for amino acids with small side chains, while ‘x’ denotes any amino acid. To note that this analysis is based only on the sequence match to the candidates considered in (Ruprecht and Kunji 2020).
